# Pheromones induce visuomotor arousal in jumping spiders

**DOI:** 10.64898/2026.10.21.760754

**Authors:** Alex M. Winsor, Guilherme F. Pagoti, Daniel Daye, Nicholas Stevenson, Ronald R. Hoy, Elizabeth M. Jakob

## Abstract

Perception in one sensory modality can be influenced by stimulation in another modality. Such cross-modal effects may arise from multiple underlying processes. For example, animals may be primed by exposure to a cue in one sensory modality and attend to congruent cues in a separate modality. In contrast, input from one sensory modality may result in a general increase in arousal, which in turn influences behavior. Jumping spiders (Salticidae) are famously visual animals but they also respond to pheromones. Here, we test whether pheromones influence visual attention in *Phidippus* jumping spiders and, if so, whether we see evidence of priming to relevant stimuli or simply an increase in arousal. In Experiment 1, male spiders exposed to female pheromones increased their movements in an arena, a sign of physiological arousal associated with mate searching, whereas females decreased their movements in response to male pheromones. In Experiment 2, freely moving male spiders exposed to female pheromones were more likely to detect and court an image of a female. In Experiment 3, male spiders were shown a series of images of conspecifics, crickets, and abstract shapes as we visualized the gaze direction of their principal eyes using a specialized eyetracker. When pheromones were present, retinal movement markedly increased, but this effect did not differ by stimulus category. Spiders showed greater retinal movement during presentations of crickets and front-facing female conspecifics than during presentations of side-view male conspecifics, regardless of pheromone exposure. In Experiment 4, spiders were shown an array of stimuli, but spiders did not preferentially look at a conspecific stimulus with pheromone exposure. In a second iteration of Experiment 4 with male spiders tested in both control and pheromone conditions, however, pheromone-exposed males looked longer at a front-facing female than at an abstract shape. In Experiment 5, spiders were presented with a salient image of a conspecific of the opposite sex. With pheromone exposure, males were more likely to divert their gaze away from the initial stimulus to an oval distractor, while females were less likely to divert their gaze than controls, mirroring sex-specific shifts in activity from Experiment 1. Increased male movement may enhance encounters, whereas reduced female movement may facilitate being found and evaluating male displays. While we found only limited evidence that spiders were primed to direct their attention toward congruent stimuli, our results show pheromones influence vision, specifically by affecting the spiders’ propensity to move their bodies and eyes.

## INTRODUCTION

In interpreting the world around them, animals often integrate information from different sensory modalities—visual, auditory, vibrational, olfactory, chemical—working in tandem (reviewed in Choi et al., 2018). Examples of multimodal integration abound across many contexts, including locating prey (Rhebergen et al., 2015), assessing the risk of predation (reviewed in Munoz and Blumstein, 2012), identifying landmarks (Schumacher et al., 2016), and evaluating courtship displays (e.g., Partan and Marler, 1999; Hebets and Papaj, 2013; Kozak and Uetz, 2016; Ligon et al., 2018; Stange et al., 2017). Multimodal integration is evident in a wide range of taxa. For example, foraging bees are better able to learn multimodal rather than unimodal cues about flower quality (Katzenberger et al., 2013), and, in humans, pairing sounds with visual stimuli improves learning in visual motion discrimination (Seitz et al., 2006).

Related to multimodal integration is cross-modal facilitation, in which stimulation in one modality improves performance in another modality (reviewed in James et al., 2022). The mechanisms underlying cross-modal facilitation can vary. For example, cross-modal cues may enhance selective attention, a process by which animals respond to specific stimuli and ignore others. Selective attention to a particular cue can be primed by exposure to a related cue, either in the same modality in a different modality. For example, humans find hidden visual targets faster in a scene after smelling a congruent odor (Seigneuric et al., 2010). Alternatively, instead of priming attention to specific stimuli, cross-modal facilitation may occur because input in one modality increases general arousal, and this increased activity and alertness improves performance on a task in another sensory modality (e.g., Olivers and Van der Burg, 2008).

Here we examine how cross-modal cues influence visual perception in jumping spiders. Jumping spiders rely on vision when hunting prey (Jackson and Pollard, 1996) and for attracting mates with elaborate courtship dances (Clark and Morjan, 2001; Lim et al., 2008; Girard et al., 2015; Glenszczyk et al., 2021). Their distributed visual system is comprised of eight eyes of two types. Three pairs of non-moving secondary eyes arrayed around the cephalothorax (head) serve as wide-field motion detectors. One pair of secondary eyes, the anterior lateral eyes (ALEs), are forward facing. The ALEs overlap in fields of view with the principal eyes, which have narrow, boomerang-shaped retinas densely packed with photoreceptors allowing for highly acute color vision and object recognition (reviewed in Harland et al., 2012; Morehouse 2020; Hill 2022; Winsor et al., 2023; Winsor et al., 2024). When the ALEs detect a stimulus, the spider directs their principal eyes toward it (Jakob et al., 2018), similar to how some vertebrates might detect a stimulus in their peripheral visual field and then move their eyes to assess the stimulus with their high-acuity foveal region. While the principal eyes have small fields of view, this constraint is mitigated because the retinas are situated at the proximal end of moveable tubes inside the spider’s cephalothorax, allowing precise targeting of gaze direction (Land, 1969).

While the visual system is often regarded as the primary sensory system in jumping spiders, other senses are also highly developed. Proprioceptive, mechanical, and chemical information are all used by salticids (reviewed in Nelson 2023). Salticids can use olfactory cues to search for host-plant refuges (Nelson et al., 2012) and prey insects (Cross and Jackson, 2010), and bitter taste to assess prey toxicity (Skow and Jakob, 2006; Vickers and Taylor, 2018; Winsor et al., 2020). Pheromones are also important: salticids can use contact chemoreception of silk draglines to ascertain the condition of nearby conspecifics (Jackson and Cooper, 1990).

Exposure to chemical cues can have cross-modal effects on visual behavior. Male jumping spiders exposed to female pheromones infused in silk show increased mate searching (Taylor 1998; Cross and Jackson, 2013). *Evarcha culicivora* jumping spiders, which prefer to feed on blood-fed mosquitoes, find hidden red prey more readily after exposure to the scent of vertebrate blood (Cross and Jackson, 2009). In a similar vein, Tedore and Johnsen (2013) found that exposure to pheromones caused male *Lyssomanes viridis* jumping spiders to court an ambiguous visual stimulus that they would have otherwise ignored or directed a threat display toward.

In this study, we test whether cross-modal pheromone cues improve visual performance, and if so, whether the improvement is due to priming of visual attention to particular stimuli or to increased arousal. First, we hypothesized that exposure to pheromones would elicit male mate search and thus lead to increased movement in an arena (Experiment 1) and increased detection and courtship of female images (Experiment 2). In our next three experiments, we used a spider eyetracker to precisely track the gaze direction of the principal eyes (Jakob et al., 2018; Canavesi et al., 2011; Bruce et al., 2021) and examine whether pheromones shift spiders’ attention toward mating-relevant visual information. While eyetracking is commonly used in humans and sometimes a handful of other vertebrate species, precise gaze tracking in invertebrates is rare but can provide novel insights (Winsor et al., 2021). We predicted that pheromone-exposed spiders would spend more time inspecting conspecific images, relative to other image types, than would control spiders (Experiment 3). We next presented spiders with an array of images and predicted that pheromone-exposed spiders would look first at the conspecific image and spend more time looking at conspecific images compared to spiders that were not exposed (Experiment 4). We also predicted that pheromone-exposed spiders would be more likely than control spiders to keep inspecting an image of a conspecific when a visual distractor appears (Experiment 5).

## METHODS

### Spider collection and housing

We used two common congeneric spiders, *Phidippus audax* and *P. princeps*, which share similar visual systems (Figure 1A), as well as life histories, including predators, hunting strategies, and mating strategies. Our choice of species for each experiment was based on their availability, which varies by season and by year. We collected spiders from several localities in western Massachusetts during the fall of 2018 (*P. princeps*) and during the fall of 2019 and 2020 (*P. audax*). Both species were captured with sweep nets and by hand in fields rich in grass and flowers (Hoefler and Jakob 2006). We also collected many *P. audax* from human-made structures, such as bridges and fences. Most individuals were adults or penultimate by the time we collected in late fall. All captured spiders were housed in individual plastic boxes (18 x 13 x 10 cm high) under a 16:8 L:D light cycle with full spectrum lighting at approximately 25°C. Once a week, we fed the spiders banded crickets (*Gryllodes sigillatus*) that were kept on a diet of high-protein dog food. We provided water *ad libitum* in vials plugged with cotton. Each spider was provided with environmental enrichment in the form of a black plastic tube, artificial leaves, and a green-painted climbing dowel (Carducci and Jakob, 2000). Male and female spider enclosures were kept on separate shelves in different areas of the laboratory to minimize visual and chemical exposure. Except where noted, spiders used in the experiments were field caught.

**Figure 1.**
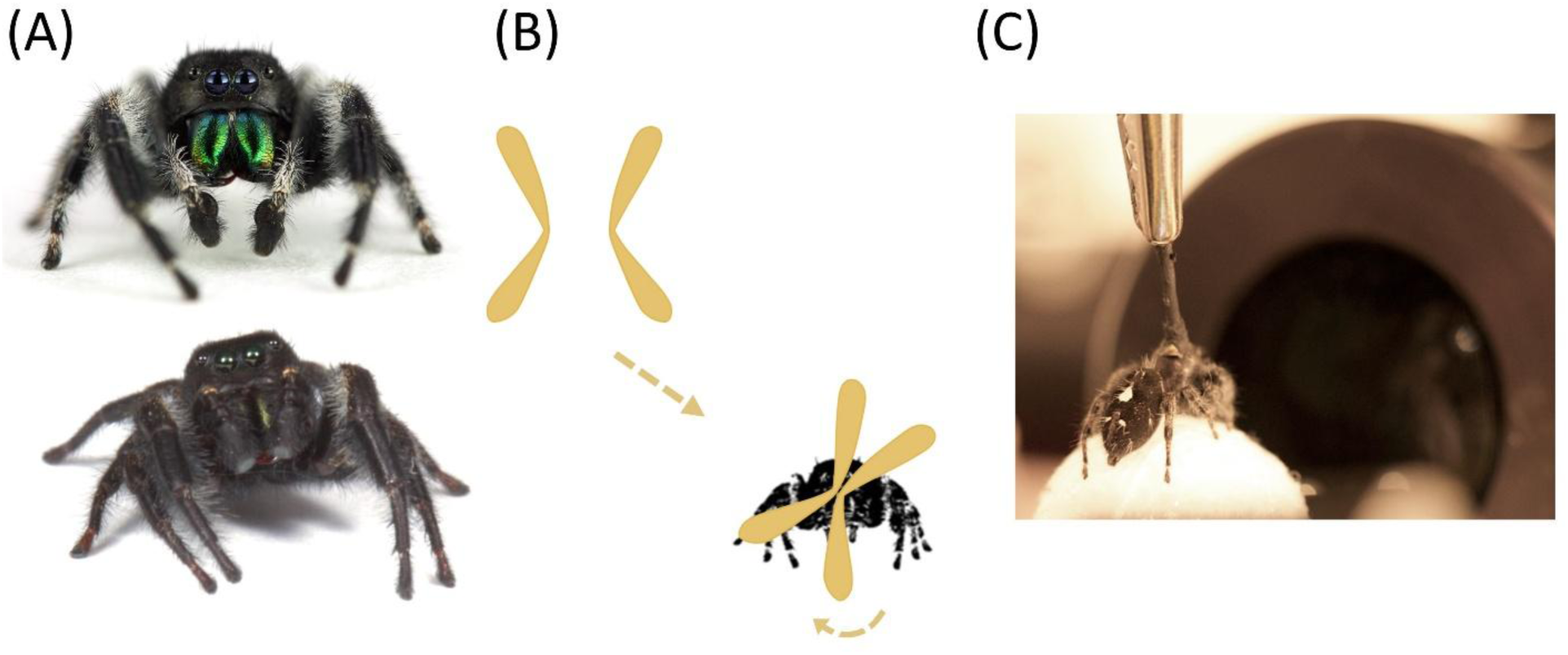
Study animals and experimental setup. (A) Anterior views of *Phidippus audax* (top) and *Phidippus princeps* (bottom), males of the two jumping spider species used for the cross-modal experiments. The large, central pair of eyes are the highly acute principal eyes, which are flanked by a pair of motion-detecting secondary eyes called the anterior lateral eyes (ALEs). (B) The vertically elongated, boomerang-shaped principal-eye retinas (yellow) come together like crosshairs to inspect objects of interest, sometimes with back-and-forth scanning and twisting motions. In this schematic, the retinas redirect toward an appearing female spider silhouette. (C) A tethered *P. audax* spider looking into the eye tracking system that allows experimenters to precisely quantify gaze direction while spiders view stimuli. The ball can be coated with pheromone cues that spiders detect with receptors on their feet and pedipalps as they walk.

### Behavioral assays overview

In Experiments 1 and 2, spiders were first exposed to pheromones or a control condition before entering a behavioral arena. To prepare the pheromone treatment, we housed each spider in separate cages (18 x 13 x 10 cm high) and fed each a cricket. One day after feeding, we transferred spiders to individual cylindrical plastic cages (15 cm diameter x 15 cm high) lined with filter paper. Spiders deposited silk lines and droppings onto the filter paper for 24 hours (Persons and Rypstra, 2001). Each spider provided only one set of cues, and each filter paper was used only once. For control stimuli, we used the same procedure but did not place a spider in the cylindrical cage. Test spiders were randomly assigned to the control or experimental group and were placed in a new, clean cylindrical cage for 10 minutes with either a clean filter paper or a pheromone paper from the opposite sex.

In Experiment 1, female and male spiders were placed in an open arena that allowed unrestricted movement. In Experiment 2, male spiders entered an arena with a long corridor terminating in a screen that displayed an image of a female.

### Eyetracker overview

In Experiments 3-5, we used a custom-built eyetracker to record the retinal movements of the principal eyes of jumping spiders (Canavesi et al., 2011). The boomerang-shaped principal-eye retinas (Figure 1B) were illuminated directly through the cephalothorax using a beam of infrared (IR) light, which is invisible to the spiders. In Experiment 3, we used an 850 nm IR light directed with a dual branch light guide equipped with a focusing lens (Thorlabs Inc., Newton, NJ, USA). In Experiments 4 and 5, we used a deeper 940 nm IR source with a focusing lens that provided improved visualization. We used an IR sensitive camera (IDS Imaging camera from Edmund Optics) to capture reflections from the retinas at 15 frames per second (fps). We presented visual stimuli in visual wavelengths to the spiders via a small projector at 30 fps (AAXA P4-X Pico). On a separate monitor, we simultaneously recorded retinal positions and the visual projection in real time using the open-source software OBS Studio (Open Broadcaster Software) at 30 fps.

We tethered spiders in the eyetracker following previously published procedures (Jakob et al., 2018; Bruce et al., 2021) (Figure 1C). Briefly, we heated a 1:1 mixture of gum rosin (Acros Organics, Fairlawn, NJ, USA) and beeswax (Stackich Inc., Troy, MI, USA) with a dental wax applicator and dipped the brush end of a microbrush (Shintop Disposable microapplicator brushes, 1.5 mm) into the melted wax. This allowed us to reversibly secure the brush to the spider’s cephalothorax. Once the wax had set, we painted the microbrush in matte black paint to reduce its reflection (Stuart Semple Black Paint, CultureHustle, Dorset, UK). The microbrush was clamped into position so that spiders could see into the eyetracker. Spiders were given a lightweight polystyrene ball to hold, which calmed them and reduced their attempts to free themselves from the tether. Spiders were allowed at least 10 min to acclimate before tests began.

To create pheromone cues for these eye tracker tests, we tethered female spiders by the cephalothorax in the same way as we tethered them for the eyetracker, described above. Tethered females were given a polystyrene ball to hold, which they could manipulate freely with their legs. Spiders “walked” on the ball while depositing dragline silk and feces for 24 hours. If pheromone balls were not used immediately after this 24-hour period, they were stored individually in sealed deli cups to reduce the diffusion of volatiles. Control balls were devoid of any draglines but were otherwise identical. Each spider was used to treat only one ball, and each ball was tested with only once.

Before each test, as described in previous studies (Jakob et al. 2018; Bruce et al. 2021), the experimenter moved a calibration square across the screen to elicit real-time saccades from the tethered spider, confirming visual responsiveness and enabling post hoc alignment of retinal and stimulus videos in Adobe Premiere Pro (v14.4.0).

## RESULTS

### EXPERIMENT 1: DOES PHEROMONE EXPOSURE INCREASE MOVEMENT?

#### Methods

We tested the effect of pheromone exposure on spider movement, an indicator of arousal. Because adult male jumping spiders engage in wide-ranging mate searching whereas adult females remain close to their silken nests (Hoefler and Jakob, 2006), we predicted that males would increase their movement in response to pheromones more than would females. We exposed field-caught adult *P. audax* spiders to opposite-sex pheromones as described previously and monitored their movements in an open-field arena.

After ten minutes in contact with clean or pheromone-infused substrate, each spider was tested in a single experimental trial in a 21.6 cm x 27.9 cm arena with walls made slick from petroleum jelly to discourage spiders from climbing out. The paper substrate in the arena was clean and was replaced between trials to remove any leftover chemical cues. We tested 30 males and 30 females once each, assigned evenly (*N* = 15) across four treatments; two control males were excluded due to escape from the arena, leaving 13 control males for analysis. Trials were recorded from directly above at 30 fps with a webcam (Logitech HD Pro Webcam C920). Trials lasted 10 minutes and we analyzed the first 5 minutes. We obtained the spiders’ pixel coordinates from the tracking software DeepLabCut (Mathis et al., 2018) and applied the distance formula between each frame. Frames with position likelihood values below 90% were rare but excluded and linearly interpolated where possible. For analysis, we built a Generalized Additive Model (GAM) with the ‘mgcv’ package (version 1.8-31; Wood, 2025) in R version 3.6.2 (R Core Team, 2019), with cumulative distance traveled as the response variable, and sex, treatment, and their interaction as explanatory variables. To measure path tortuosity, a measure of trajectory complexity, we divided the total path length by the net displacement. The initial and final positions were defined as the average x, y position across the first and last 30 frames, respectively, to improve the accuracy of net displacement estimates. We used a Wilcoxon rank-sum test in R to analyze path tortuosity. To measure speed, we averaged the distance traveled per minute. For analysis, we built a GLM with gamma-distributed errors and used Holm pairwise comparisons for the groups of interest.

#### Results

After pheromone exposure, males showed greater spatial exploration than control males, whereas females showed reduced spatial exploration relative to control females (Supplementary Figure 1; Supplementary Video 1). Total distance traveled differed among groups, with the highest values observed for pheromone-exposed males and control females relative to the other groups (Figure 2A; Supplementary Table 1; pheromone males: 78.1 ± 0.16 cm, control females: 77.6 ± 0.17 cm, pheromone females: 57.0 ± 0.14 cm, and control males: 52.0 ± 0.17 cm, using a conversion factor of 1 cm ≈ 26.4 pixels; GAM: *P* < 0.05). Path tortuosity was highest for pheromone-exposed males but lowest for control female spiders (Figure 2B; Wilcoxon rank-sum test: P = 0.04). For movement rate, there was a significant interaction between sex and treatment (GLM: β_pheromone*Male_ = 0.5110 ± 0.2511, *t* = 2.035*, P* = 0.047), with pheromone-exposed males moving faster on average than control males (923.9 ± 87.7 vs. 642.9 ± 131.8 px/min), and pheromone-exposed females moving more slowly on average than control females (692.1 ± 83.1 vs. 889.2 ± 100.8 px/min). From the pairwise comparisons, this interaction was most strongly driven by males (Holm contrast between control and pheromone males: -0.344 ± 0.181, *df* = 54, *t* = -1.901, *P* = 0.0626). Overall, pheromone exposure increased total distance traveled in males but decreased it in females.

**Figure 2.**
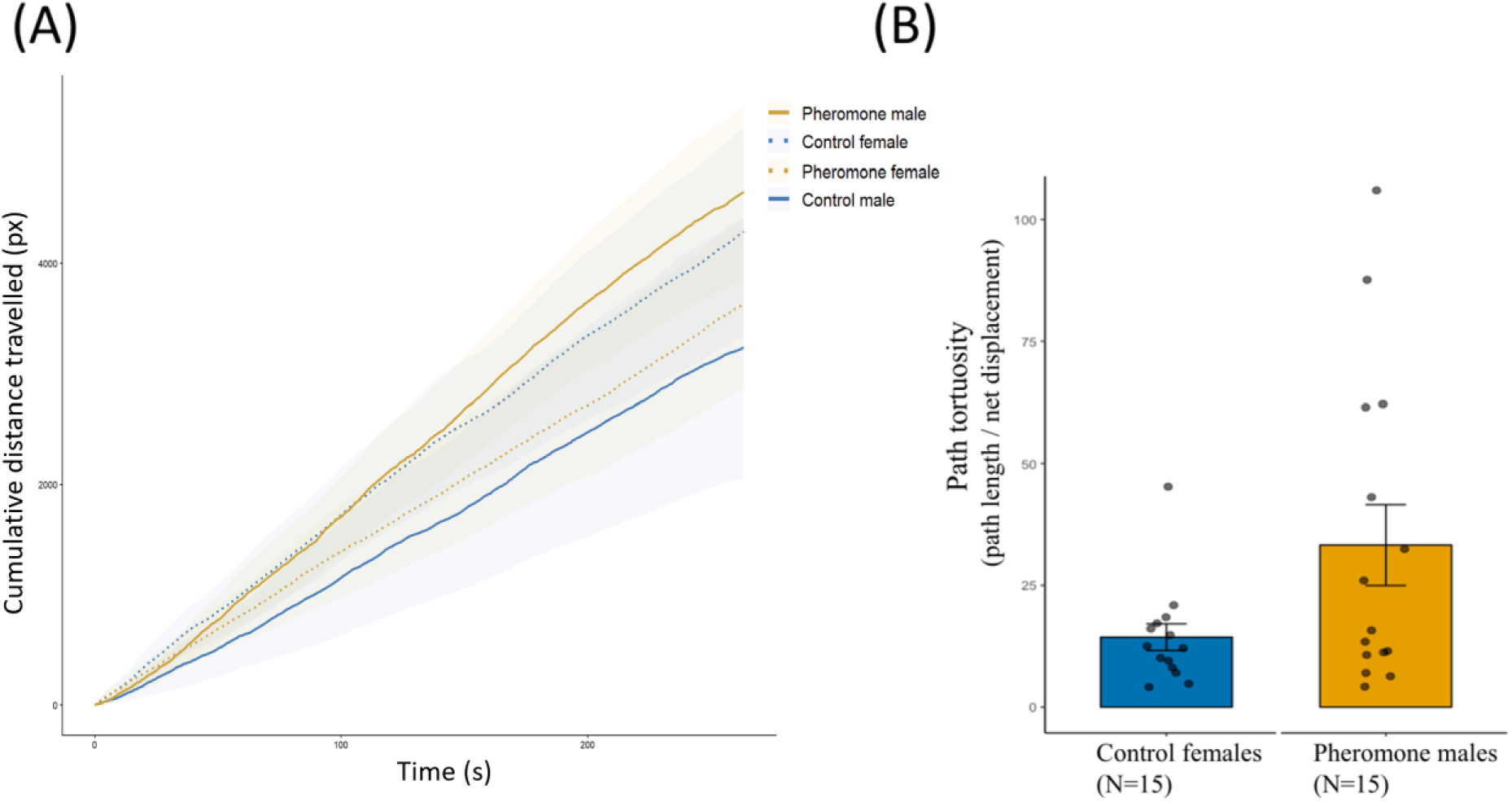
Locomotor activity increased for pheromone-exposed males. (A) The average total distance traveled (i.e., the total pixel-distance traveled averaged across individuals within a group) was highest for pheromone-exposed males. In contrast, after pheromone exposure, females had a lower distance traveled. (B) Path tortuosity, which explains the twistiness of the locomote path, was highest for pheromone-exposed males and lowest for control females—the two groups with highest overall movement rates. Bars show means and their standard errors, with individual spider values depicted as jittered points. Together, these results suggest that after pheromone exposure, males engage in local search while females remain in a similar spot throughout the trial.

### EXPERIMENT 2: DOES PHEROMONE EXPOSURE INCREASE DETECTION AND COURTSHIP OF FEMALE IMAGES?

#### Methods

If pheromones facilitate visual performance, exposure to female pheromones should increase the propensity of male spiders to detect and court images of prospective mates. Here we tested only male *P. aduax*. In pilot trials, all males courted live females regardless of pheromone exposure, indicating a strong baseline courtship response, so we presented a more ambiguous cue: a video of a photo of a female that moved back and forth. Thus, no cues from local motion were present, but the movement of the photo kept the attention of the spider. We presented the video on an Apple iPod (generation 5; Apple Inc., Cupertino, CA) positioned against on one the short walls of a rectangular arena made of foam board (15 cm x 9.5 cm x 11 cm height). Trials were conducted in a room under full spectrum lights (Philips 20W 24 in T12 Daylight).

Pheromone and clean filter papers were prepared, and males were exposed as in Experiment 1. After pheromone exposure, a spider was transferred to the clean testing arena and left to acclimate for three minutes under a small transparent condiment cup (3 x 2 cm) positioned at the long end of the arena opposite from the iPod. We then started the stimulus video and released the spider. Trials were recorded with a webcam (Logitech HD Pro Webcam C920) at 30 fps positioned 20 cm above the arena on a tripod. We scored the latency for males to detect the female (defined as orientation of its body to direct its forward-facing eyes to the image), and the times at which it started and ended courtship (defined as raising the front pair of legs and moving side-to-side while oriented to the image). Trials lasted 3 minutes. We tested 40 spiders divided equally (*N* = 20) between the treatment and the control groups.

Because this experiment involved simple comparisons between two independent treatment groups, we used nonparametric Wilcoxon rank-sum tests in R to assess the effect of pheromone exposure on the latency to orient to the female image, the number of courtship bouts directed to the female image, and the total duration of all courtship bouts.

#### Results

While all spiders in both groups eventually detected the female image, pheromone-exposed males detected the female image faster (Figure 3A; Wilcoxon rank sum test, *W* = 276.5, *P* = 0.015). Pheromone-exposed males also exhibited significantly more bouts of courtship during the trial than did non-exposed animals (Figure 3B; *W* = 111.5, *P* = 0.012). Although pheromone-exposed males showed a greater total courtship duration across the entire trial (*W* = 105.0, *P* = 0.006), this effect was driven by whether males courted at all. When analyses were limited to males that exhibited at least one courtship bout, total courtship duration did not differ between treatments (*W* = 13, *P* = 0.267). Thus, pheromone exposure increased both the likelihood and number of bouts of courtship behavior.

**Figure 3.**
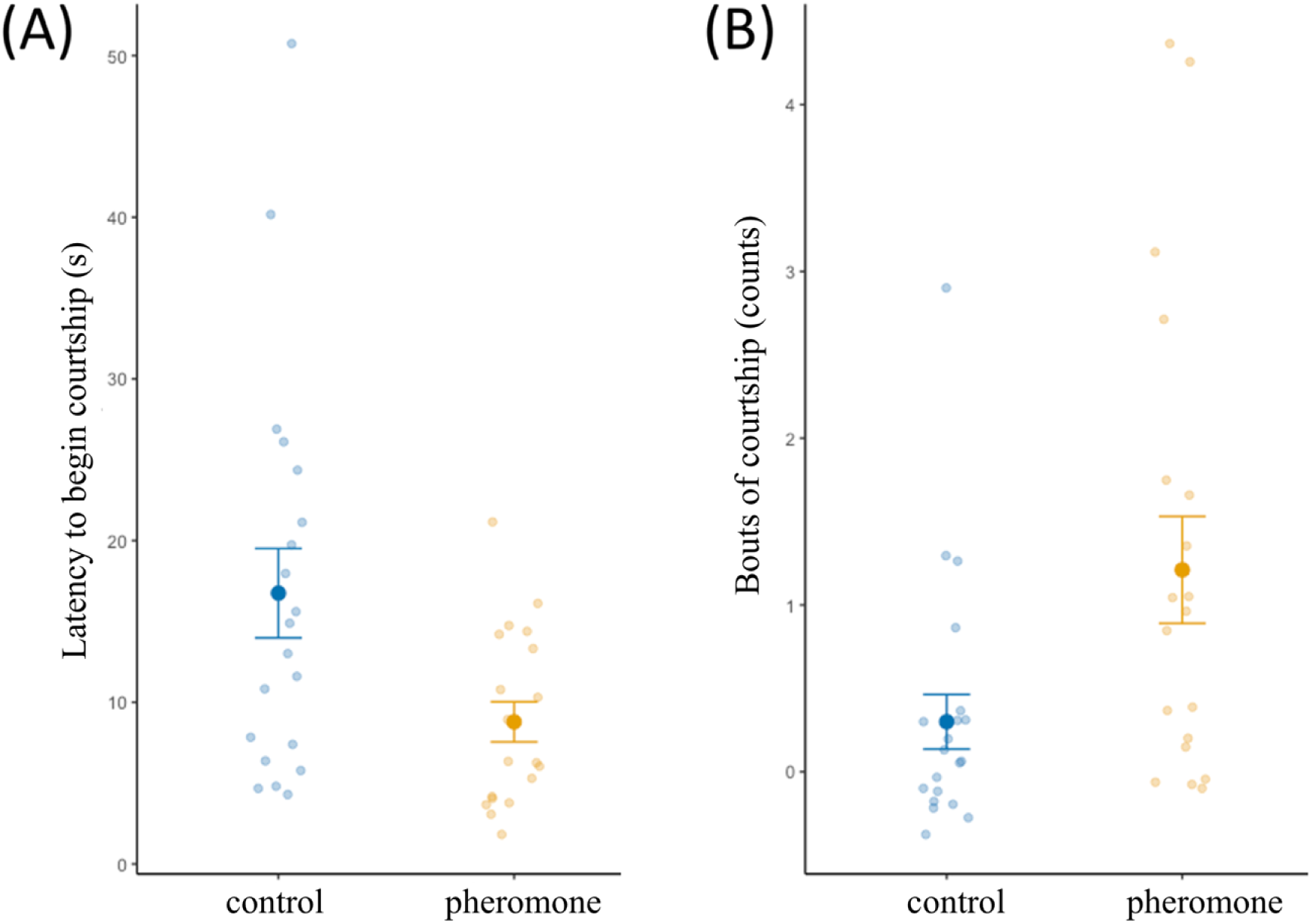
Pheromone exposure increased male detection and courtship of female images. (A) Control males had a longer latency to detect the female image, compared to pheromone exposed males. (B) Pheromone-exposed males exhibited more bouts of courtship in response to the image. Means are depicted as a large point with standard error bars, while smaller, jittered points represent individual spider values. The increased number of courtship bouts is consistent with pheromone-exposed males entering a reproduction-like behavioral state.

### EXPERIMENT 3: DOES PHEROMONE EXPOSURE INFLUENCE ATTENTION TO SEQUENTIALLY PRESENTED IMAGES?

#### Methods

In the previous experiment, we found that freely moving males were significantly more likely to court female images if they were exposed to pheromones. Here we tested whether pheromone exposure influenced male gaze direction in the eyetracker (its design is described under General Methods). For this, we showed spiders a sequence of images: anterior and lateral views of both male and female conspecifics, a cricket silhouette, and a shaded oval (Figure 4). We predicted that pheromone-exposed males would look longer at images of spiders than at other images and would look longer at female images than would control males. Due to limited availability of field-caught spiders, we used 12 *P. princeps* tested twice under each condition, with and without pheromones.

**Figure 4.**
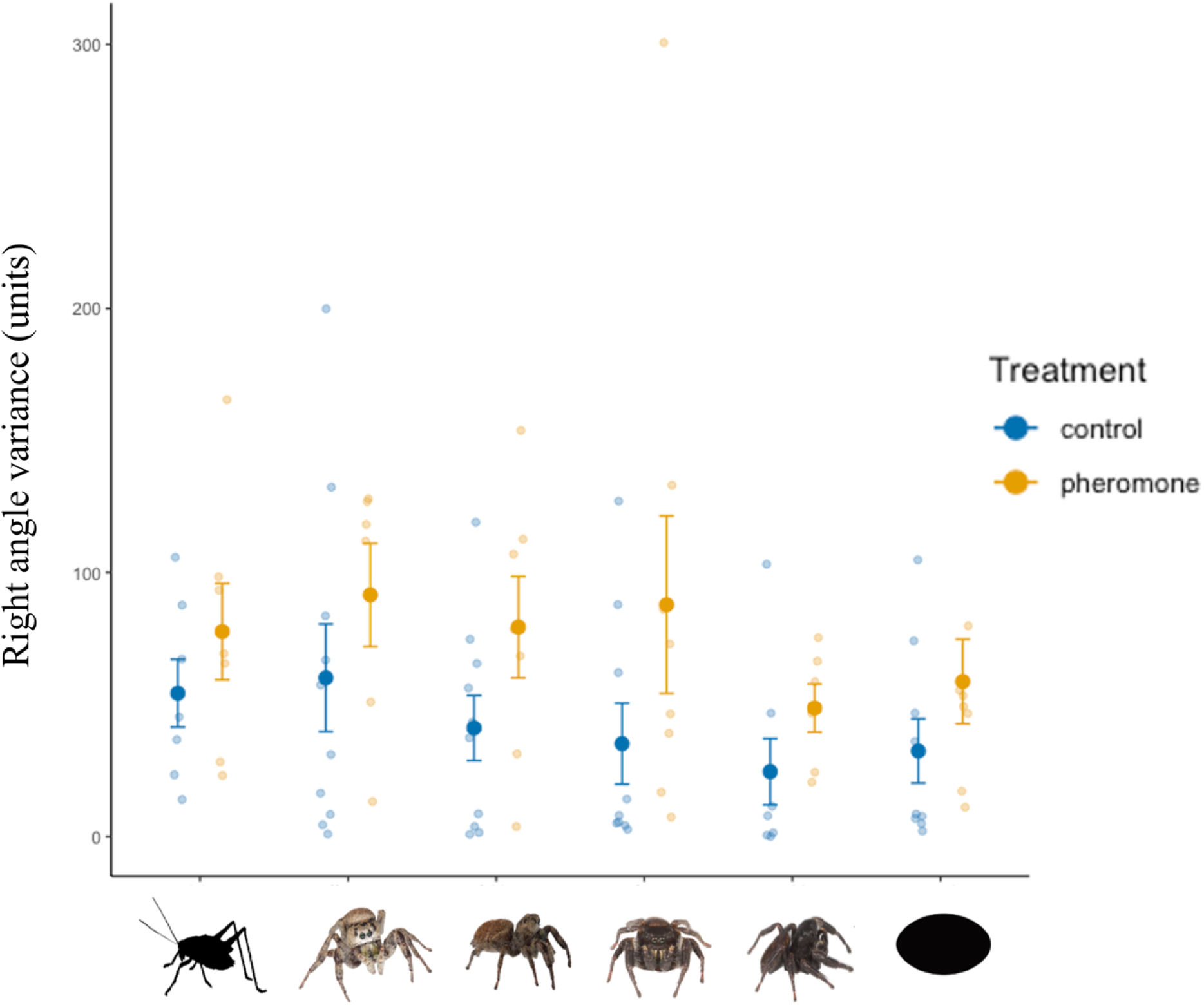
Pheromone exposure elicits an overall increase in retinal activity. Shown on the x axis are the stimuli that were presented in random order to the spiders, while the average amount of retinal torsion is displayed on the y axis. Means are depicted as a large point with standard error bars, while smaller, jittered points represent individual averaged trial values. Each spider was tested twice with the stimulus set, one time for each treatment. Regardless of stimulus type, the retinas always twisted and moved more with pheromone exposure, suggesting arousal rather than priming for a congruent visual cue.

For testing, we tethered each male spider in the eyetracker, provided them with either a pheromone or control ball chosen at random, and used the calibration routine described previously to ensure that spiders were aligned such that all presented stimuli appeared in the center of the visual field. For each trial, spiders were given a 20-min pre-trial acclimation period with a blank grey screen while holding the ball. Next, spiders were shown a randomized series of six static images, each with an equal number of pixels. Each image was shown for 60 seconds, and each image presentation was followed by a five-minute break with a dark screen. Next, the ball was removed and replaced with one from the opposite treatment. After a 20-min break while holding the new ball, spiders were shown the images again in a new random order.

For each image presentation, we measured retinal behavior in four ways. First, because spiders actively explore images that they are interested in, we measured the distance that retinas moved during the course of a 60-second image presentation. Second, spiders usually examine stimuli via scanning, or tilting their retinas back and forth (Land 1969). To quantify the degree of scanning for each stimulus presentation, we measured the change in the angle of the retina from one frame to the next, and then calculated the variance in angle changes; a higher variance reflected more scanning behavior. Third, when spiders are exploring an image of interest, they place the high-acuity inner “elbows” of the two boomerang-shaped retinas close together (Figure 1B), so we measured the inter-retinal distance. Finally, we also measured the distance of each “elbow” to the center of the stimulus.

To extract retinal position, we overlaid the stimulus and retinal movement videos as described previously and exported them using Adobe Media Encoder as JPEG image stacks. We tracked the position of each retina frame by frame using the plugin MTrackJ in the program Fiji (v2.0.0-rc-69/1.52p, Schindelin et al., 2012). In each frame, we manually annotated the center point of the “elbow” and the top of each retina. These measures together enabled us to calculate retinal angle. Because *Phidippus* retinas tend to exhibit torsion in synchrony (Jakob et al., 2018), we arbitrarily calculated angles of the right retinas only. We also annotated the position of the stimulus center. The treatment of each spider was unknown to the observers annotating the image stacks, and therefore data were collected blind. We then calculated changes from one frame to the next (as in Jakob et al. 2018).

All data processing and analyses were done in R. Generalized linear mixed effects models (GLMMs) were run using the function ‘glmer’ from the package ‘lme4’ (version 1.1-21, Bates et al., 2015). To examine if there were differences in retinal movement, torsion, inter-retinal distance, and stimulus-retinal distance, we ran GLMMs with gamma-distributed errors and a log link function, with the treatment of the spider (control or pheromone) and stimulus type (6 levels) treated as factors, and the spider ID treated as a random effect. Non-significant parameters were dropped from the final model. Model fit and residual diagnostics were assessed with the ’DHARMa’ package (Hartig and Hartig, 2017).

#### Results

We found that pheromone exposure influenced three of our four measures of retinal movement, but stimulus order and the interaction between stimulus type and pheromone treatment were not significant for any measure (Supplementary Table 2). Retinas of pheromone-exposed spiders demonstrated significantly greater variance in retinal angle relative to controls, indicating increased scanning behavior (Figure 4; GLMM: β_pher_ = 1.000 ± 0.200, *t* = 4.992, *P* < 0.001). Pheromone-treated spiders also moved their retinas greater distances than those of control spiders (GLMM: β_pher_ = 0.672 ± 0.246, *t* = 2.734, *P* = 0.006) and exhibited higher rates of retinal speed (Supplementary Figure 2). Moreover, with pheromone exposure, the distance between retinal centers was larger (GLMM: β_pher_ = 0.185 ± 0.063, *t* = 2.943, *P* = 0.003), consistent with reduced retinal convergence during visual exploration. In contrast, we found no significant differences in the average distance of the retinas relative to the stimulus center for any independent variable, including pheromone treatment (GLMM: *P* > 0.05). Regardless of pheromone exposure, we detected differences in spiders’ responses to various stimuli. In our reduced GLMM model, retinal torsion was greater during presentations of crickets and front-facing female conspecifics relative to the oval stimulus intercept, and pairwise comparisons further showed greater activity for both stimuli than for the side-view male conspecific (Tukey contrast between cricket and male side: 1.087 ± 0.341, *t* = 3.184, *P* = 0.018; female front vs male side: 1.012 ± 0.321, *t* = 3.152, *P* = 0.020). Thus, pheromone exposure increased retinal activity, and retinal activity also varied among stimulus images, but pheromone exposure did not alter stimulus-specific responses.

### EXPERIMENT 4: DOES PHEROMONE EXPOSURE INFLUENCE ATTENTION TO IMAGES PRESENTED IN AN ARRAY?

#### Methods

In the previous experiment, where images were presented in succession, pheromones significantly increased overall retinal movement but did not differentially bias responses toward specific stimulus types, possibly because tethered spiders just oriented to each image as it appeared. Males also exhibited greater retinal activity in response to the front-facing female image compared to views of a same-sex conspecific. In Experiment 4, we presented different images simultaneously in an array, thus requiring spiders to choose some images over others. We predicted that spiders exposed to pheromones of the opposite sex would be more likely to direct their gaze at an image of an opposite-sex conspecific compared to competing images.

In this experiment, we used male and female *P. audax*. Pheromone and control balls were created with identical methods from Experiment 3. Spiders were aligned in the eye tracker as above and given a randomly selected pheromone or control ball to walk on. To begin the trial, spiders tracked a moving target around the screen with saccades and scanning fixations. When the target was at the center of the screen and the spider was scanning it, a hotkey was pressed by the experimenter, revealing a four-way array of images equidistant from the target’s center and whose positions were randomly assigned.

We ran two variations of this experiment with different groups of spiders. For the first, we tested wild-caught spiders (*N* = 58) once each, randomly assigned to control or pheromone-treatment groups. For the second variation, we tested lab-reared spiders (*N* = 54) twice each in an attempt to look for individual differences in behavior, once with and once without pheromones in random order, with tests at least two days apart. To rear spiders, we separated spiderlings after they emerged from their egg sacs, placed them into individual 16 oz deli cups and fed them crickets, flour beetle larvae (*Tribolium* sp.), and fruit flies (*Drosophila melanogaster*) three times weekly. After the final molt, all laboratory conditions were identical to captive wild-caught spiders.

Stimulus arrays and presentation parameters differed between the two iterations of Experiment 4. For the first iteration, the array consisted of an anterior view of an opposite-sex conspecific, a cricket, an oval, and a cross (Supplementary Figure 3A). The array was presented for five minutes and shook side-to-side (∼1 Hz with a 56-px peak-to-peak displacement, 4 px per frame) for two seconds every fifteen seconds to maintain the spiders’ attention. For the second iteration, the array consisted of an anterior and lateral view of an opposite-sex conspecific, a cricket, and a cross (Supplementary Figure 3B). Here we included a lateral view of a conspecific because we reasoned that this view might be more readily encountered by males during mate searching. Because we had observed in the first variation that spiders’ attention flagged over the course of the trial, we reduced the presentation time to one minute, with two seconds of shaking every five seconds.

Overlays of the retinas and stimuli were made in Adobe Premiere Pro 2019, which were aligned using an established calibration protocol (Bruce et al., 2021; Jakob et al., 2018). Each treatment was exported as an .mp4 file from Adobe Media Encoder and scored via video playback. For scoring, observers blind to the pheromone treatment used ethogram event logging software (BORIS; Friard and Gamba, 2016) to track where spiders where looking. We scored a spider as looking at a stimulus when the high-acuity central “elbows” of both principal-eye retinas were brought together over an image. Each such instance was defined as a bout of looking. We recorded the first stimulus chosen, number and duration of bouts, and total looking time toward each stimulus.

We ran generalized linear models (GLMs) and generalized linear mixed-effects models (GLMMs) using the function ‘glmer’ from the package ‘lme4’ (Bates et al. 2015) and the function ‘glmmTMB’ from the package ‘glmmTMB’ (Brooks et al., 2017) in R. For our first analysis, we used a GLM (iteration 1 of the experiment) and a GLMM (iteration 2) with binomially distributed errors and a logit link function to test whether the first stimulus selected in the array was a conspecific image or not, which would suggest that information gathered by the secondary eyes was used to immediately direct the principal eyes to particular images. Our second analysis tested whether time spent looking at conspecific images changed with pheromone exposure, sex, or an interaction between them, using a GLM (iteration 1) and GLMM (iteration 2) with gamma-distributed errors and a log link function. In another analysis for iteration 2, we sought to test if a larger proportion of time was spent looking at spider images compared to the other two images. Since the array consisted of two views of a conspecific image, rather than aggregating total looking time for spider, we quantified how viewing time was distributed between spider and non-spider images using a GLMM with beta-distributed errors and a logit link function. For this analysis, we analyzed only males (*N* = 32), each tested twice, as pheromone-mediated effects of visual search were expected to be strongest in males, and we used the proportion of time these males spent looking at conspecifics relative to the other images as the response variable. Model fit and residual diagnostics for all analyses were assessed using the ‘DHARMa’ package (Hartig and Hartig, 2017).

### Results

Similar to the previous experiment, we found no evidence that pheromone exposure altered stimulus-specific attention in either iteration of Experiment 4. In the first iteration, using wild-caught spiders, there was no significant difference in the first stimulus selected or the amount of looking time to the conspecific (Figure 5; Supplementary Table 3A; Supplementary Videos 2-3; *P* > 0.05) across the fixed effects of treatment, sex, or their interaction in GLM models. During the five-minute presentation, spiders viewed the array for a total average of 30.5 seconds, the remaining time of which was spent viewing blank parts of the screen.

**Figure 5.**
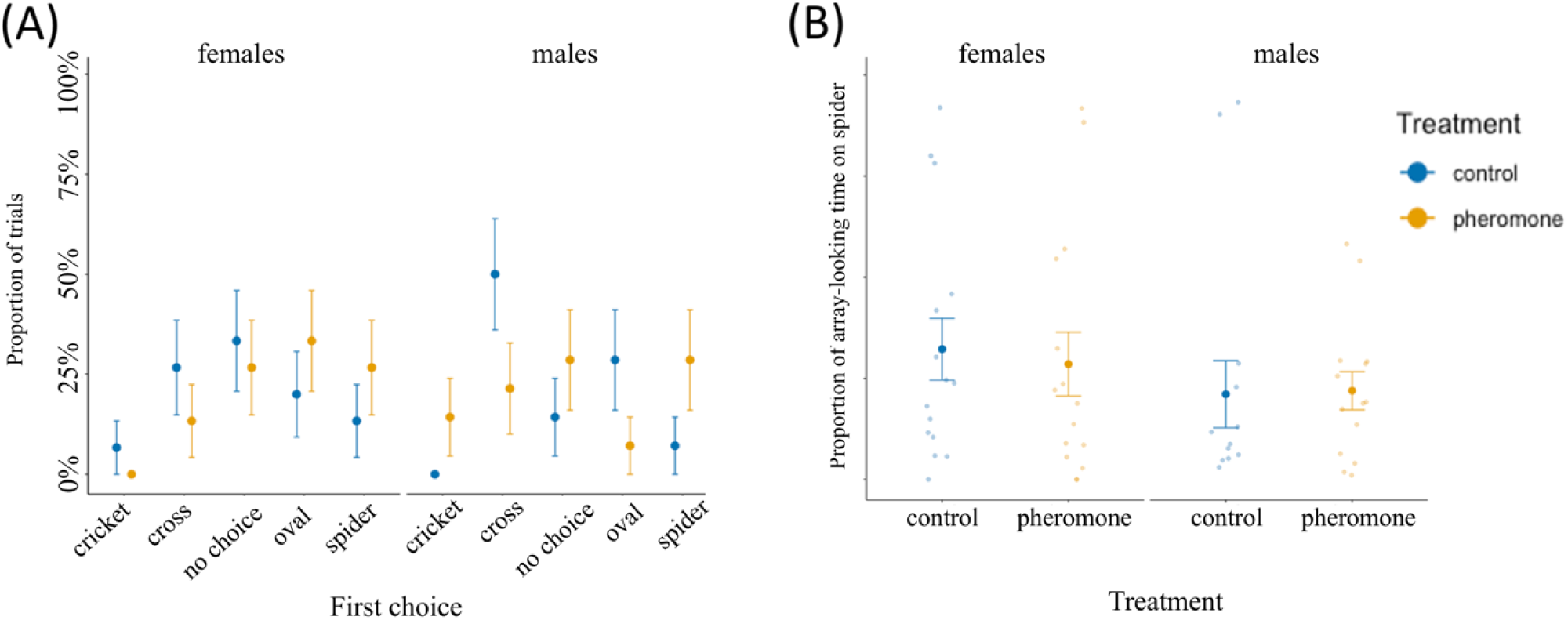
There was no difference in first choice or proportion of time spent looking at the spider image. (A) The probability of selecting a particular stimulus first was not different across sex or treatment groups. The N/A choice occurred when spiders immediately disengaged with the array as it appeared, and thus no choice was made. (B) The proportion of time spent looking at the spider relative to the rest of the array was not different across sex or treatment. Jittered points represent the response of individual spiders, with bars denoting the standard error of the mean across conditions. Plotted is the first iteration of the experiment with wild-caught spiders, but lab-reared spiders showed similar response patterns.

In the second iteration of Experiment 4 with male lab-reared spiders, treatment did not significantly affect the first stimulus selected or the proportion of looking time directed toward both conspecific views compared to the rest of the array (GLMMs: *P* > 0.05). When looking time was instead analyzed separately for each stimulus using a GLMM with gamma-distributed errors, pheromone-exposed spiders spent more time looking at the front-facing female than at the abstract cross (Supplementary Table 3B; Tukey contrast between front-facing female and cross: 3.87 ± 1.95, *df* = 244, *t* = 2.69, *P* = 0.038). During the one-minute presentation, spiders viewed the array for a total average of 8.1 seconds, which was comparable to the first iteration when controlling for differences in stimulus presentation times (13.5% of total looking time from lab-reared spiders compared to 10.1% of total looking time from wild-caught spiders). For spiders that were tested twice, one time in each treatment, variation was dominated by trial-to-trial variability (overdispersion = 2.69) rather than among-individual differences (GLMM random intercept variance = 0.23).

### EXPERIMENT 5: DOES THE PRESENCE OF PHEROMONES DECREASE ATTENTION TO A DSITRACTING STIMULUS?

#### Methods

In the previous experiment, spiders did not preferentially reorient toward images of conspecifics following pheromone exposure; however, we next asked whether pheromones influenced the maintenance of attention once a spider was already fixating on a conspecific image. We presented spiders (*N* = 54) with a modified distractor protocol used in earlier experiments to test the control of attention of the principal eyes by the secondary eyes (Bruce et al., 2021). First, a primary visual stimulus was presented to a spider in the eyetracker. Once the spider was scanning the primary stimulus with back-and-forth, torsional retinal motions of its principal eyes, a distractor appeared in the field of view of the secondary eyes (i.e., the ALEs). We positioned the distractor outside of the field of view of the principal eyes as they examined the stimulus, but in an area of the screen that they could be redirected to see. Thus, a spider would detect the distractor with the ALEs and then redirect—or not redirect—their high-acuity principal eyes to examine the distractor. Here our goal was to test whether spiders were less likely to be distracted from looking at a conspecific image when they were exposed to pheromones.

We used the same spiders from the second iteration of Experiment 4 and conducted Experiment 5 directly afterwards (with half of the individuals receiving Experiment 5 first). Spiders remained tethered in the eyetracker and were given a 5-minute break with a blank screen before the initiation of Experiment 5. Spiders, still holding their control or pheromone-infused ball, were shown a primary stimulus of an opposite-sex conspecific *P. audax*. Once their retinas began to scan the object, a hotkey was pressed by the experimenter which revealed a pixel-matched oval distractor visible only in the field of view of the secondary eyes. Spiders could either ignore the distractor and continue to look at the primary stimulus or direct their retinas to the distractor (scored as a binary choice, as no spiders looked away from all stimuli). Five spiders were excluded as they never looked at the initial stimulus, for a final sample size of *N* = 49.

A generalized linear mixed effects model (GLMM) was built using the function ‘glmer’ from the package ‘lme4’ (Bates et al. 2015) in R. We used binomial distributed errors with a logit link function, assigning order, treatment, sex, and the interaction between treatment and sex treated as fixed effects, and spider ID treated as a random effect. Model fit and residual diagnostics were assessed with the ’DHARMa’ package (Hartig and Hartig, 2017).

#### Results

In the final experiment, we tested whether spiders looked away from an image of an opposite-sex conspecific and toward a suddenly appearing oval. There was a significant interaction between sex and pheromone exposure: when pheromones were present, males were more likely to look at the distractor, whereas females were less likely (GLMM: β_sex*pheromone_ = 2.51 ± 1.012, *z* = 2.48, *P* = 0.013; Figure 6; Supplementary Table 4; Supplementary Videos 4-5), compared to when pheromones were not at all present, mirroring results from the first experiment. That is, with pheromone exposure, males increase their general activity while females decrease theirs.

**Figure 6.**
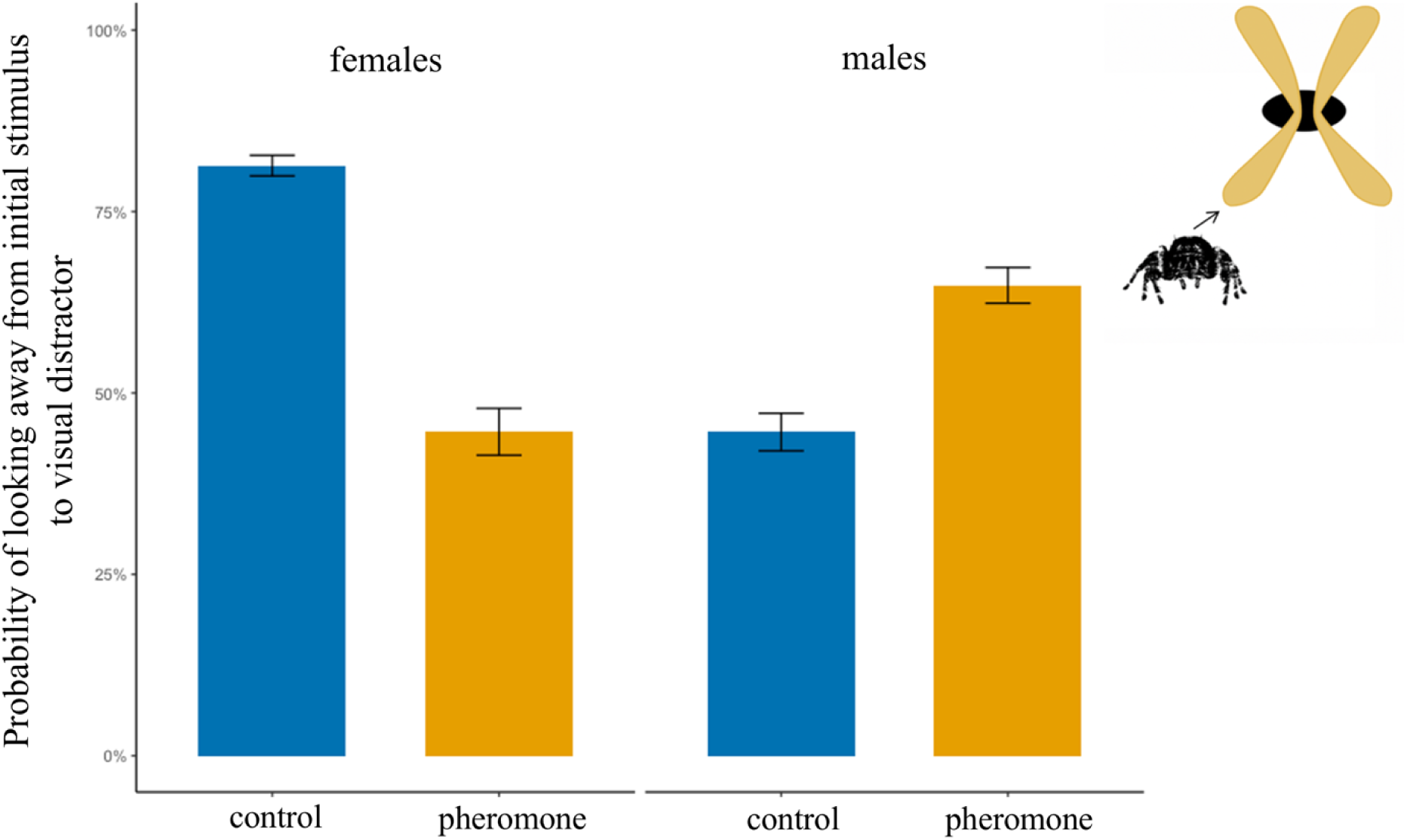
With pheromone exposure, males were more distractable while females were less distractable. Spiders were initially focused on an opposite-sex conspecific image but could choose to redirect their gaze to an appearing oval distractor (inset, upper-right corner). Bars show GLMM model-estimated probabilities of orienting to an appearing distractor, with bars representing the standard error of the mean. There was an interactive effect between treatment and sex, mirroring results from Experiment 1. With pheromone exposure, males increased their activity, while females decreased theirs.

## DISCUSSION

We demonstrate that principal eyes of male jumping spiders move significantly more when spiders are exposed to pheromone cues. However, we found limited evidence that spiders are more likely to direct their gaze toward particular stimuli that are congruent with the cross-modal pheromone cues. Instead, overall, our evidence suggests that pheromones increase arousal rather than priming attention to particular cues.

Our first two experiments examined behavior of freely moving spiders in arenas. Males exposed to female pheromones were more active, whereas exposure to male pheromones decreased female movement (Experiment 1). These results align with spider behavior in the field: adult female *Phidippus* generally remain relatively stationary as they inspect male courtship displays, as well as when they built silken nests where they rest at night and during inclement weather, and where they lay their eggs. In contrast, males roam considerable distances in search of females and often share the nests of females rather than building their own (Jackson, 1986; Hoefler, 2007; Hoefler and Jakob, 2006). These results are in accord with the idea that physiological arousal increases when males are exposed to pheromones.

In Experiment 2, we tested the effect of pheromone exposure on the attention of freely moving males to an image of a female. Males that were exposed to female pheromones detected the image more rapidly and were more likely to court the image compared to males that were not exposed to pheromones. These results are similar to that of Tedore and Johnsen (2013), who found that pheromone exposure increases the likelihood that jumping spiders interpret visual images as potential mates. However, this experiment cannot show whether spiders were more likely to find the female image simply because they were in a higher state of arousal, or because they were primed to be more attentive to female presence.

In Experiments 3 and 4, we address the issue of visual attention more directly by examining eye movement. We found that eye movements increased significantly when male spiders were exposed to pheromones, suggesting that pheromone exposure alters retinal sampling and, consequently, the visual information acquired. However, contrary to predictions, we found no evidence that spiders exposed to pheromones were more likely than controls to attend to images of conspecifics compared to other images, whether images were presented sequentially to the spider (Experiment 3) or in an array (Experiment 4). Nonetheless, spiders showed overall greater retinal movement during presentations of crickets and front-facing female conspecifics in Experiment 3, and pheromone-exposed males in the second iteration of Experiment 4 spent more time looking at the front-facing female than at a shape control. This suggests that spiders, at least as demonstrated by these tasks, were not consistently primed by pheromones to look for relevant images, despite some evidence that frontal views of a female conspecific were especially salient to male spiders. Two primary explanations may account for this finding. First, an increase in eye actively may simply indicate an increased general arousal, rather than more directed selective attention such as priming. Higher arousal may still be adaptive, as an increase in retinal sampling would increase the probability that a potential mate in the visual field would be detected against the background, which might explain our findings in Experiment 2. Second, it may be that our static images were not recognized by the spiders, whereas images with realistic biological motion (De Agrò et al. 2021; De Agrò et al., 2025) might have been classified appropriately. However, this second explanation seems less likely, as untethered spiders respond appropriately to both nonmoving three-dimensional objects (Experiment 2; Rößler et al., 2021) and fairly crude two-dimensional images of spiders (Drees 1952). Thus, frontal conspecific views may provide especially salient visual information during mate assessment.

In Experiment 5, we found that females were more likely to keep looking at a front-facing conspecific image when presented with a distracting visual stimulus when they were exposed to pheromones compared to when they were not. In contrast, pheromone-exposed males were more likely to be distracted from looking at a female image by a suddenly appearing stimulus. These sex differences may be understood in light of Experiment 1 and the natural mating behavior of jumping spiders. During courtship, male jumping spiders initiate theatric courtship displays, often misdirecting courtship to heterospecifics (Taylor et al., 2017) and sometimes to inanimate objects. In contrast, receptive females generally orient toward a courting male and hold relatively still to carefully assess his display (Echeverri et al., 2017). A general increase in arousal, in both body and eye movements, might allow males to discover and redirect their behavior to court more potential mates and ward off more approaching male competitors, while in turn, decreased arousal or more focused attention on the nuances of the display might allow females to be found easier and help them choose the best mate. In Experiment 5, females often continued to inspect the male image using torsional retinal movements in spite of a distractor, suggesting that they may be evaluating it in some way.

Our results suggest that in these species, pheromone cues might regulate behavioral state, rather than recruiting higher cognitive processes such as search image formation. This pattern mirrors other work showing that predator sounds increase eye movements without preferentially directing gaze toward a potentially threating visual stimulus (Winsor et al., companion submission). Therefore, shifts in visual attention are likely to be a product of the general modulation of arousal, possibly mediated by mechanisms such as neuromodulation. Serotonin and octopamine levels, which have been shown to modulate visual processing (Cheng and Frye, 2020), and dopamine levels, which have been shown to regulate behavioral responsiveness (van Swinderen et al., 2008), might rapidly fluctuate after exposure to cross-modal cues and initiate a shift in behavioral state and underlying sensory processing. In some contexts, neuromodulation might be more computationally efficient and a better use of resources than higher cognitive processes. Future research measuring endogenous fluctuations of bioamines in the brain or hemolymph, such as with high-performance liquid chromatography, would be informative to this end. It is perhaps underappreciated that animals likely employ a combination of lower and higher-level cognitive tasks to meet selective demands imposed by their ecology. Future research should also seek to integrate the role of specific object features across modalities, behavioral state, and integration of sensory inputs in the brain during cognitive tasks.

Increasing retinal sampling to acquire additional information at critical times is likely very important for animals, as is ignoring irrelevant information. Given the ubiquity of eye movements across the animal kingdom, recently found even in insects (Fenk et al., 2022), how sequences of eye movements are used across taxa to extract information from a scene is an important area of future research. Eye-tracking studies can offer unparalleled insights into animal cognition and behavior (Winsor et al., 2021; Billington et al., 2020). Work precisely measuring gaze direction with non-primate animals has burgeoned in recent years, with studies in birds (Tyrrell et al., 2014; Yorzinski et al., 2017), flies (Fenk et al., 2022), spiders (Bruce et al., 2021), cephalopods (Feord et al., 2020), and mice (Michaiel et al., 2020), all revealing interesting insights. Given the independent evolution of eye movements in several lineages, first for gaze stabilization, and later for foveation of a scene (Land, 2019), looking around more might have been a precursor to looking around *for something*.

## Acknowledgments

We thank Ethan Clotfelter, Sarah Partan, Lisa Mangiamele, Laela Sayigh, Renae Brodie, Jeff Podos, Luke Remage-Healy, and Paul Katz for providing comments on drafts of the manuscript. We also thank Agustin Plasencia and Alyssa Wilkinson for independently rescoring the array data, and Meagan Gustafson and Zachary Fisher for independently rescoring the distractor data as a reliability check. This research was funded by NSF IOS 1656714 to E.M.J. and R.R.H.

## SUPPLEMENTARY INFORMATION

**Supplementary Figure 1.**
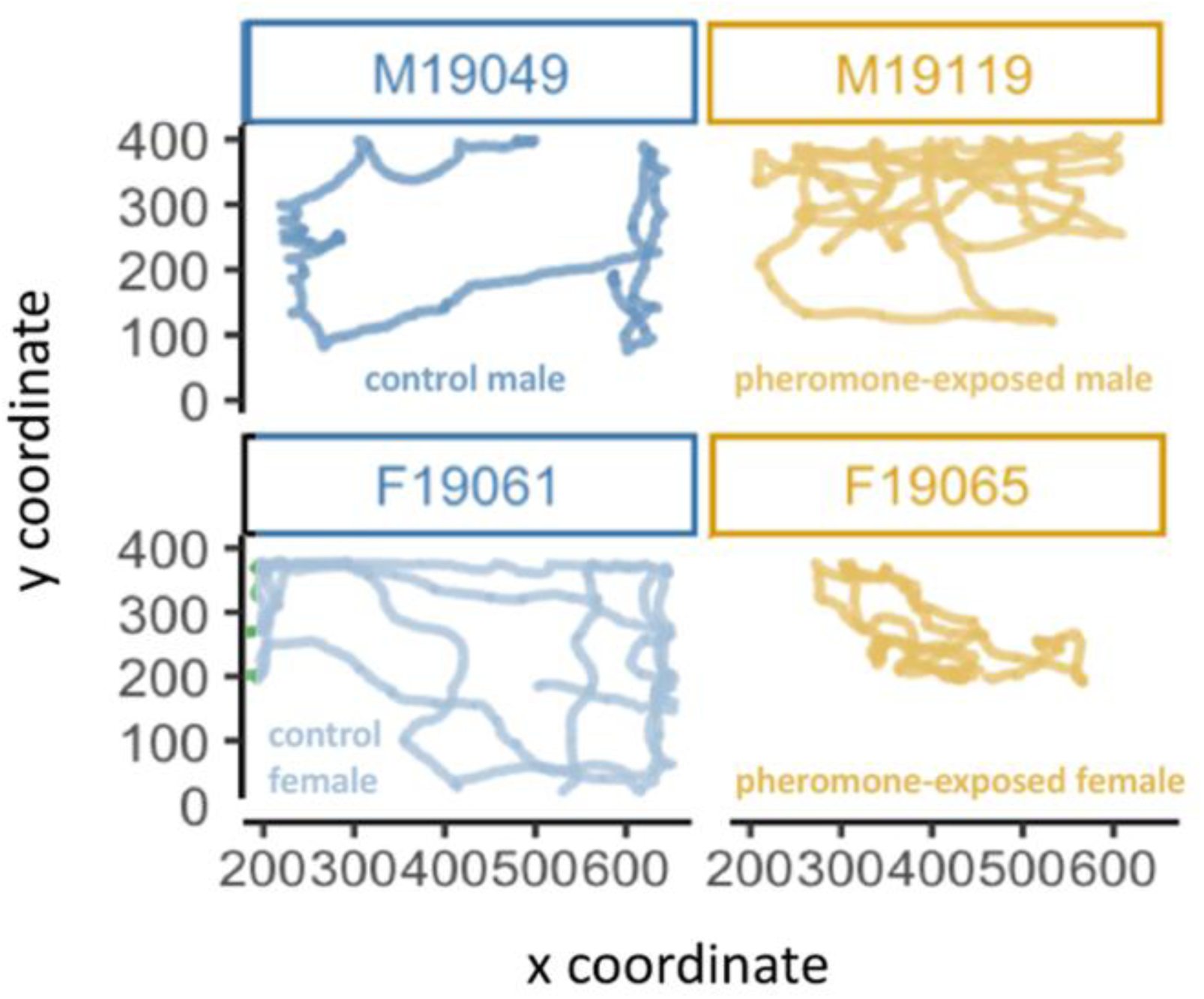
Representative traces of the locomotion path used by a spider from each treatment. The left and right columns display control and pheromone-exposed spiders, respectively, while the top row shows control and pheromone-exposed males, and the bottom row shows control and pheromone-exposed females. Darker shading indicates greater overlap of traces and therefore greater spatial occupancy in a particular location. All spiders started the trial in the center of the arena. After pheromone exposure, males moved around more. By contrast, females, which typically engaged in wall-walking behavior in the control condition, moved around less when exposed to pheromones.

**Supplementary Figure 2.**
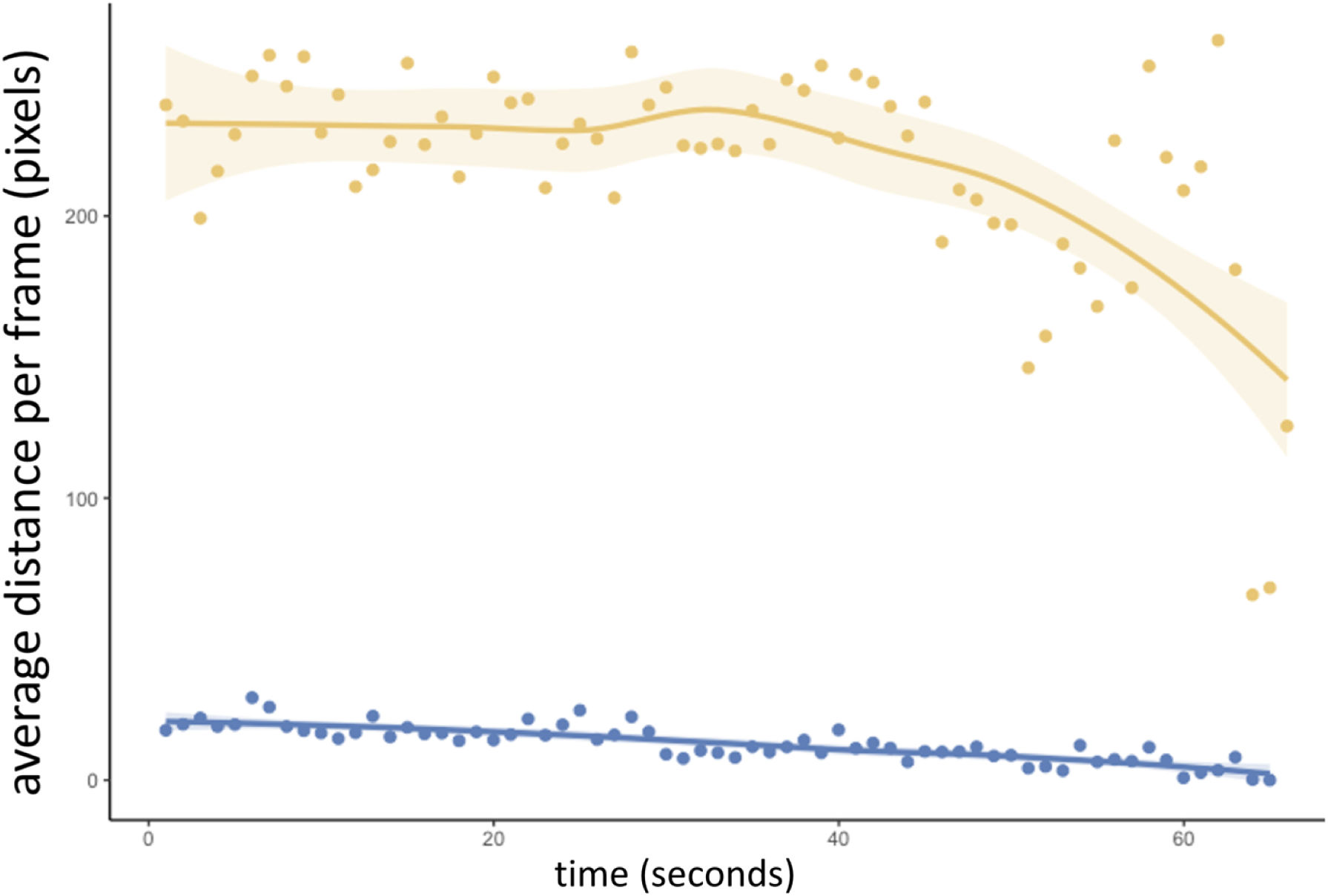
Retinal speed increases with pheromone exposure. Pheromone-treated values are shown in gold, while control values are show in dark blue. Speed was calculated as the Euclidean distance between retinal positions in consecutive frames and plotted over time after down sampling from 15 to 1 fps. Curves show mean distance traveled averaged across all stimulus presentations and individuals, with points indicating averages at each time bin and shaded areas indicating ± SEM. Retinal speeds are far higher in the pheromone condition, even toward the end of the trial when interest wanes.

**Supplementary Figure 3.**
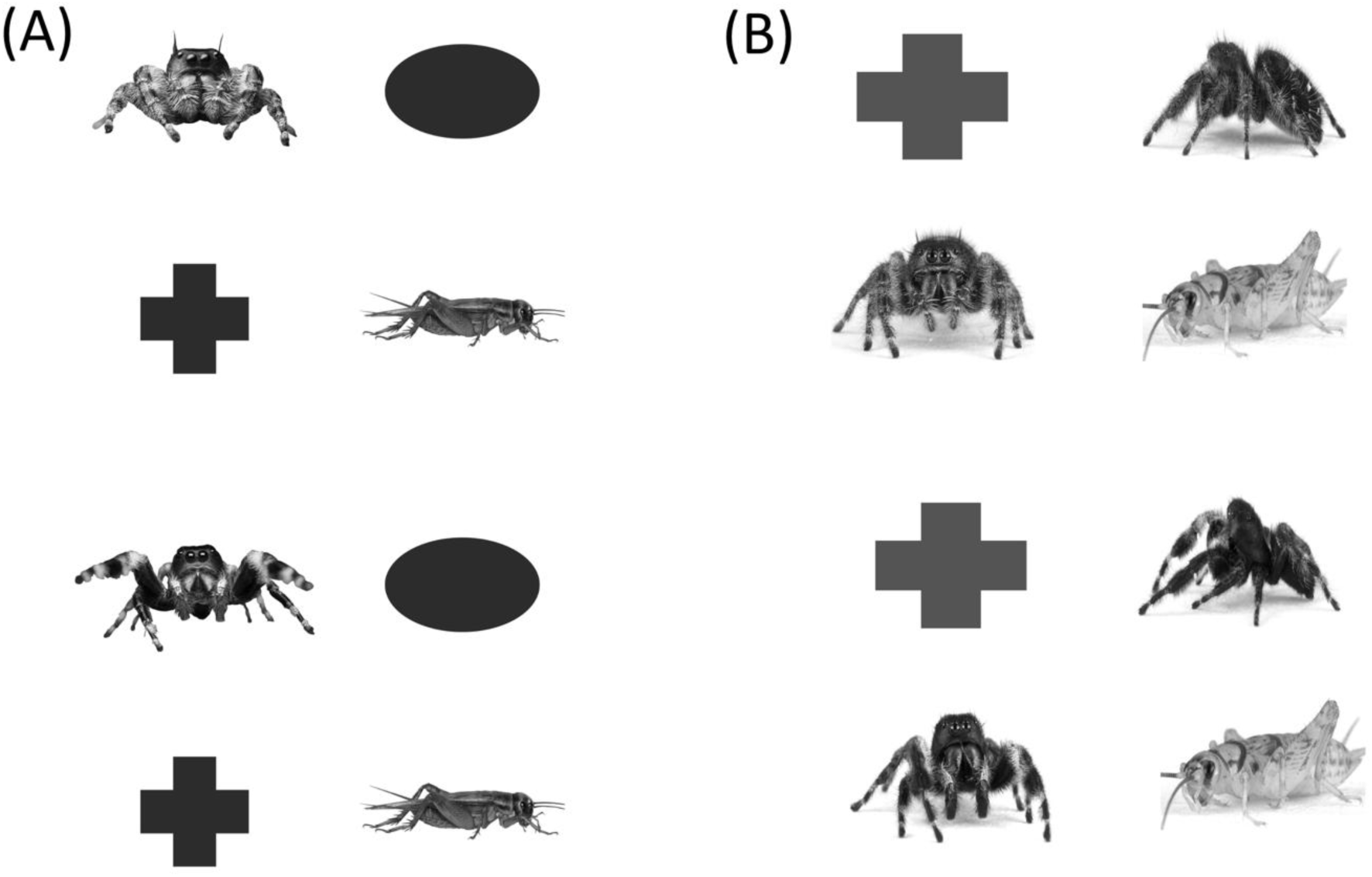
Four-way array stimuli used in Experiment 4. (A) Array used in the first iteration with wild-caught spiders tested once, which consisted of an opposite-sex conspecific, cricket, cross, oval. The array that was shown to males is on top, while the array that was shown to females is on the bottom. (B) Array used in the second iteration with lab-reared spiders tested twice, which consisted of an anterior view of an opposite-sex conspecific, a lateral view of an opposite-sex conspecific, a cricket, and a cross. The array shown to males is on top, while the array shown to females is on the bottom. Stimulus position was randomly shuffled on every trial. The array design drew on approaches used in the human psychology literature. Visual search, the process of selecting relevant items from a scene (Wolfe 2010; Spaethe et al., 2010), might have involved rapid first choices mediated by the secondary eyes (consistent with parallel-like search), or sequential inspection with prolonged viewing by the principal eyes (consistent with parallel-like search), but we found no strong evidence for either.

**Supplementary Table 1.**
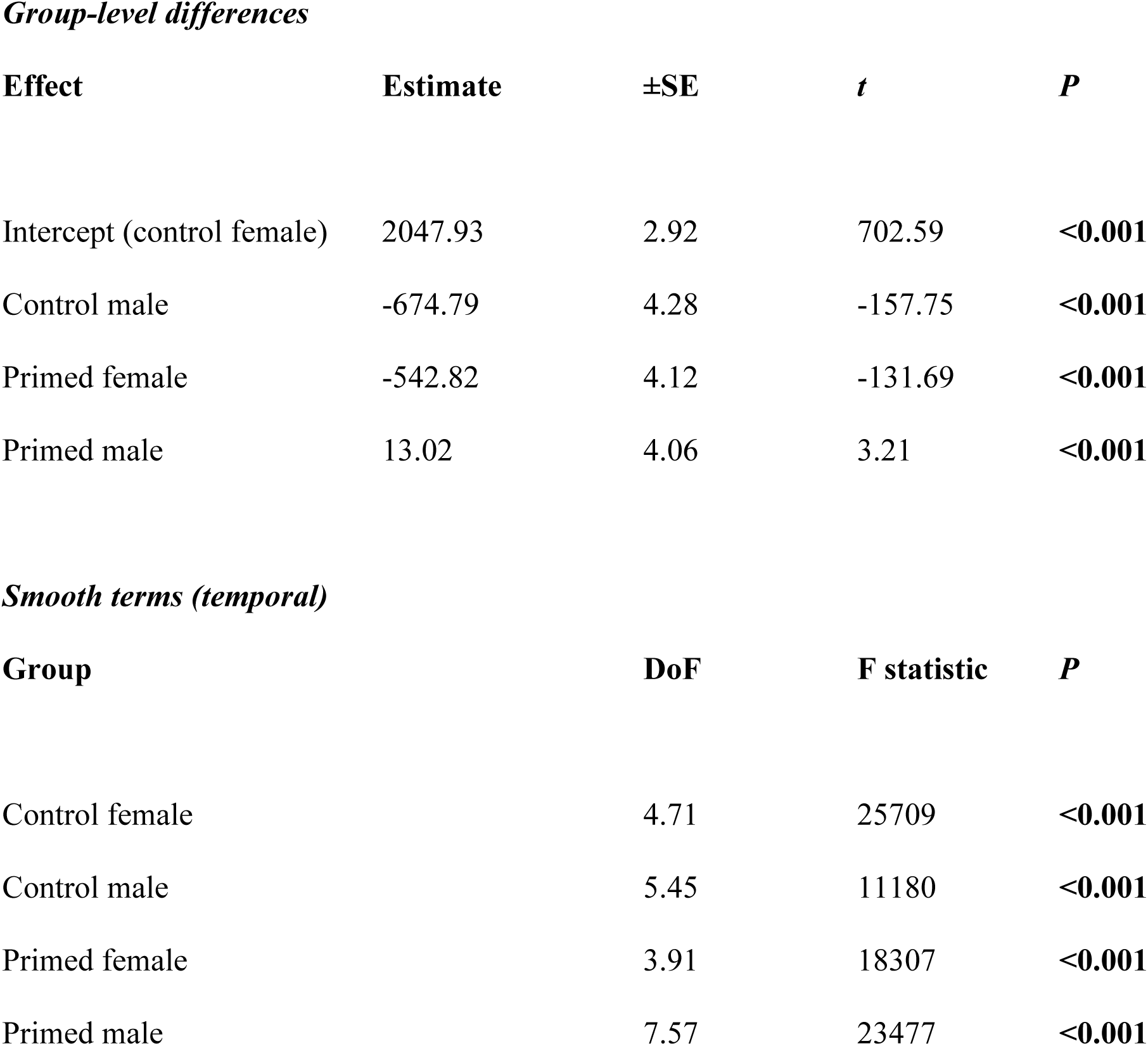
GAM model estimates (±SE) of total distance traveled across treatments. Values are relative to the intercept, which represents control females. The *t* and *P* values are indicated for each fixed effect. Estimated degrees of freedom (DoF), *F*, and *P* values are reported for group-specific smooth terms describing nonlinear changes in movement over time. Bolded significant P values indicate groups that differ significantly in overall movement magnitude or temporal trajectory relative to the reference group.

| **Effect** | **Estimate** | **±SE** | ***t*** | ***P*** |
| --- | --- | --- | --- | --- |
| Intercept (control female) | 2047.93 | 2.92 | 702.59 | **<0.001** |
| Control male | -674.79 | 4.28 | -157.75 | **<0.001** |
| Primed female | -542.82 | 4.12 | -131.69 | **<0.001** |
| Primed male | 13.02 | 4.06 | 3.21 | **<0.001** |

**Supplementary Table 1. GAM model estimates (±SE) of total distance traveled across**
| **Group** | **DoF** | **F statistic** | ***P*** |
| --- | --- | --- | --- |
| Control female | 4.71 | 25709 | **<0.001** |
| Control male | 5.45 | 11180 | **<0.001** |
| Primed female | 3.91 | 18307 | **<0.001** |
| Primed male | 7.57 | 23477 | **<0.001** |

**Supplementary Table 2.** GLMM model estimates (±SE) of right retinal angle variance. Values are relative to the intercept which represents control spiders that were shown an oval. T and P values are indicated for each fixed effect. The bolded significant P values show pheromone-exposed spiders exhibited significantly more retinal torsion than control spiders. For the reduced model, where the interaction terms are dropped, the front-facing female stimulus (ffront) rises to significance at P < 0.05.

| Factor | Estimate | ±SE | *t* | P |
| --- | --- | --- | --- | --- |
| Intercept (oval control) | 3.05895 | 0.37342 | - | - |
| treatment is pheromone | 0.92553 | 0.41928 | 2.207 | **0.0273\*** |
| cricket:pheromone | -0.55850 | 0.61214 | -0.912 | 0.3616 |
| ffront:pheromone | 0.08116 | 0.62028 | 0.131 | 0.8959 |
| fside:pheromone | 0.42378 | 0.60191 | 0.704 | 0.4814 |
| mfront:pheromone | 0.34509 | 0.59712 | 0.578 | 0.5633 |
| mside:pheromone | 0.24706 | 0.62534 | 0.395 | 0.6928 |
| stimulus cricket | 1.01800 | 0.43622 | 2.334 | **0.0196\*** |
| stimulus ffront | 0.66657 | 0.39250 | 1.698 | 0.0895 |
| stimulus fside | 0.23897 | 0.40110 | 0.596 | 0.5513 |
| stimulus mfront | 0.08523 | 0.41309 | 0.206 | 0.8365 |
| stimulus mside | -0.45080 | 0.41782 | -1.079 | 0.2806 |

**Supplementary Table 3.**
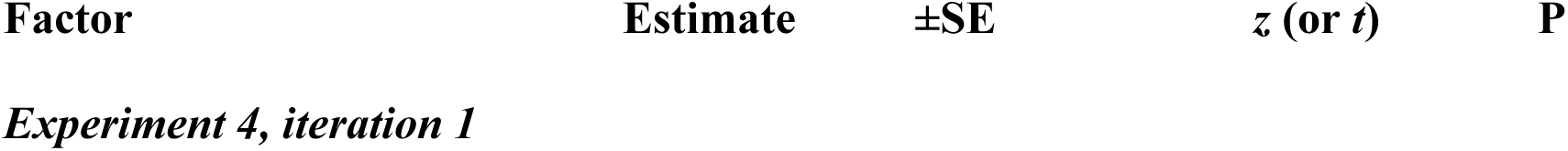
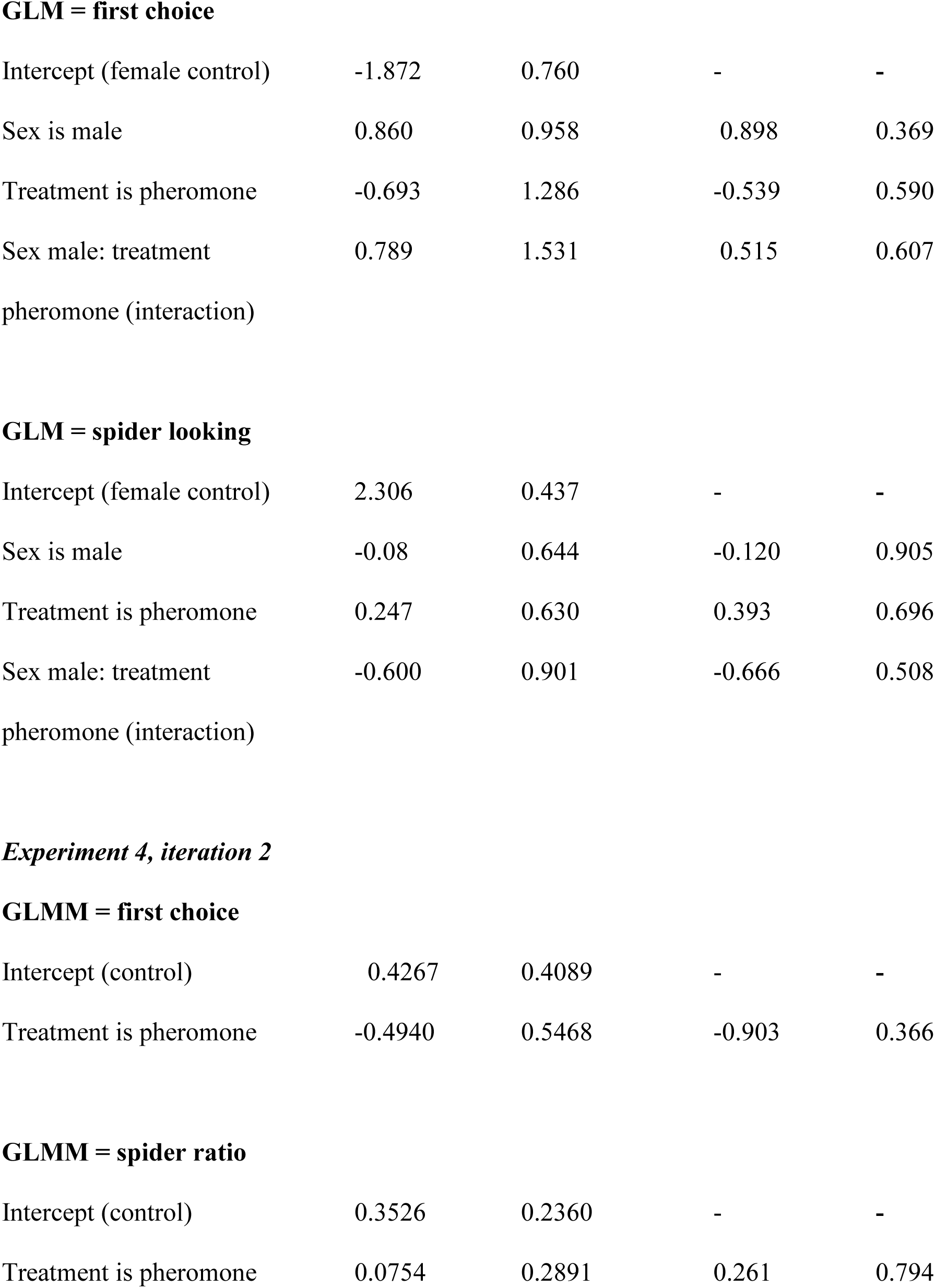
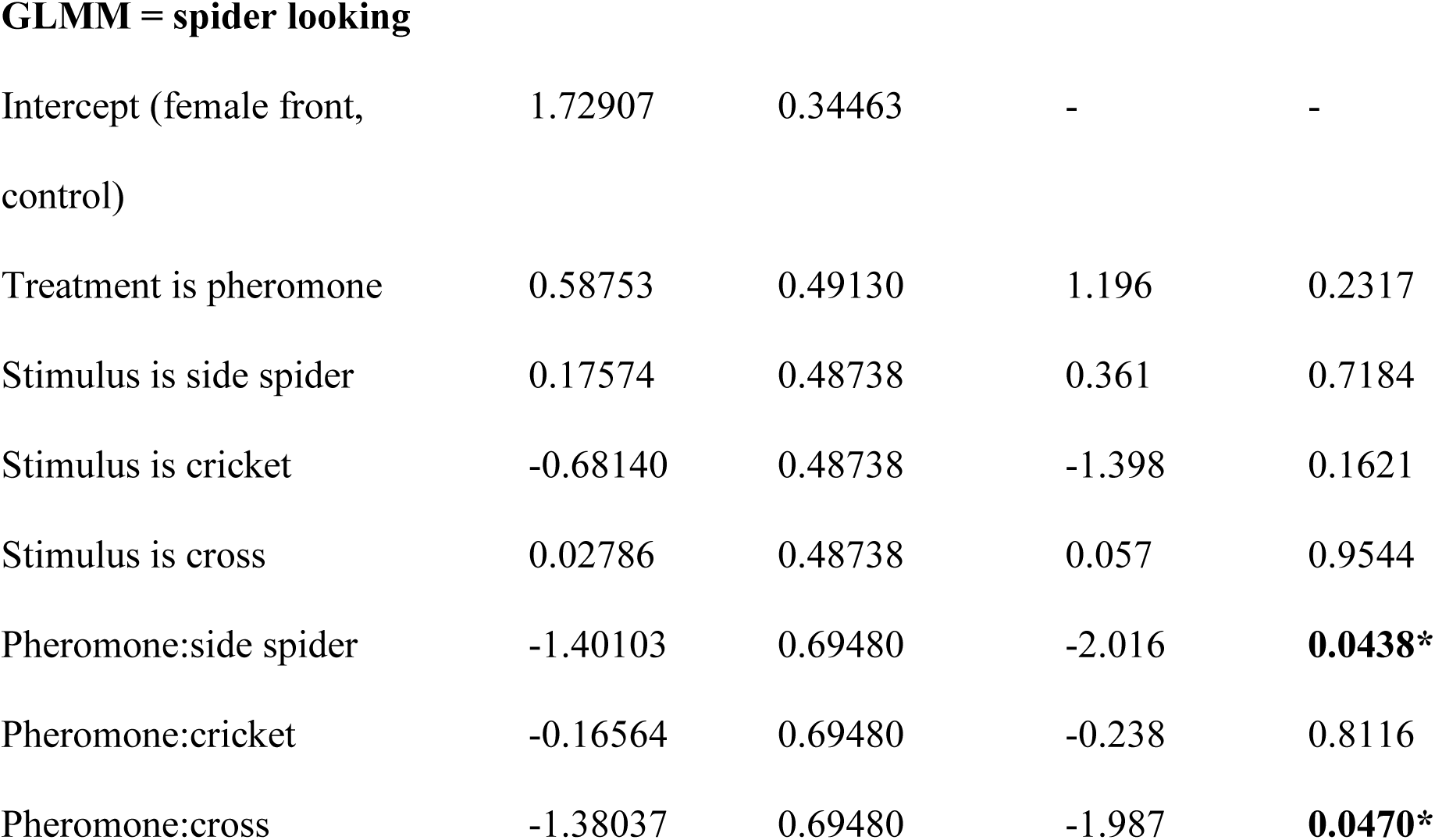
Model estimates (±SE) of the probability of a spider looking first to the image of a conspecific in an array, and model estimates (±SE) of looking time, and proportion of total looking time, to conspecific images. Values are relative to the intercept which represents a control (control female in the first iteration, control viewing female front in the second iteration). Z and P values are indicated for each fixed effect. The lack of significant effects of pheromone exposure on first choice, looking time, or the proportion of looking directed toward conspecific views suggests that pheromones did not broadly increase selective attention to conspecific stimuli. However, in the second iteration, pheromone-exposed males showed greater looking toward the forward-facing female conspecific view relative to comparison stimuli, suggesting some stimulus-specific selectivity within the pheromone condition. The first iteration of Experiment 4 is displayed on top (GLMs with binomial and gamma-distributed errors), while the second iteration with only males (GLMMs with binomial, beta, and gamma-distributed errors along with a spider-level random effect for repeated measures) is shown on the bottom.

| Factor | Estimate | ±SE | *z* (or *t*) | P |
| --- | --- | --- | --- | --- |
| *Experiment 4, iteration 1* |  |  |  |  |

**GLM = first choice**

|  |  |  |  |  |
| --- | --- | --- | --- | --- |
| Intercept (female control) | -1.872 | 0.760 | - | - |
| Sex is male | 0.860 | 0.958 | 0.898 | 0.369 |
| Treatment is pheromone | -0.693 | 1.286 | -0.539 | 0.590 |
| Sex male: treatment<br>pheromone (interaction) | 0.789 | 1.531 | 0.515 | 0.607 |

**GLM = spider looking**

|  |  |  |  |  |
| --- | --- | --- | --- | --- |
| Intercept (female control) | 2.306 | 0.437 | - | - |
| Sex is male | -0.08 | 0.644 | -0.120 | 0.905 |
| Treatment is pheromone | 0.247 | 0.630 | 0.393 | 0.696 |
| Sex male: treatment<br>pheromone (interaction) | -0.600 | 0.901 | -0.666 | 0.508 |
| ***Experiment 4, iteration 2*****GLMM = first choice** |  |  |  |  |
| Intercept (control) | 0.4267 | 0.4089 | - | - |
| Treatment is pheromone | -0.4940 | 0.5468 | -0.903 | 0.366 |

**GLMM = spider ratio**

|  |  |  |  |  |
| --- | --- | --- | --- | --- |
| Intercept (control) | 0.3526 | 0.2360 | - | - |
| Treatment is pheromone | 0.0754 | 0.2891 | 0.261 | 0.794 |

**GLMM = spider looking**

**Supplementary Table 3. Model estimates ( ± SE) of the probability of a spider looking first to the image of a conspecific in an array, and model estimates ( ± SE) of looking time, and proportion of total looking time, to conspecific images.**
|  |  |  |  |  |
| --- | --- | --- | --- | --- |
| Intercept (female front, control) | 1.72907 | 0.34463 | - | - |
| Treatment is pheromone | 0.58753 | 0.49130 | 1.196 | 0.2317 |
| Stimulus is side spider | 0.17574 | 0.48738 | 0.361 | 0.7184 |
| Stimulus is cricket | -0.68140 | 0.48738 | -1.398 | 0.1621 |
| Stimulus is cross | 0.02786 | 0.48738 | 0.057 | 0.9544 |
| Pheromone:side spider | -1.40103 | 0.69480 | -2.016 | **0.0438\*** |
| Pheromone:cricket | -0.16564 | 0.69480 | -0.238 | 0.8116 |
| Pheromone:cross | -1.38037 | 0.69480 | -1.987 | **0.0470\*** |

**Supplementary Table 4.** GLMM model estimates (±SE) of the probability of a spider disengaging with a conspecific stimulus to look at an oval distractor. Values are relative to the intercept which represents a control female. Z and P values are indicated for each fixed effect. The bolded significant P values show that there was interaction between sex and treatment, such that their response was opposite relative to each other.

| Factor | Estimate | ±SE | z | P |
| --- | --- | --- | --- | --- |
| Intercept (female control) | 1.472 | 0.615 | - | - |
| Sex is male | -1.688 | 0.756 | -2.234 | **0.026 \*** |
| Treatment is pheromone | -1.687 | 0.778 | -2.168 | **0.032 \*** |
| Sex male: treatment<br>pheromone (interaction) | 2.514 | 1.015 | 2.477 | **0.013 \*** |

## SUPPLEMENTARY VIDEOS

https://drive.google.com/file/d/1PdPJ1StHaiWr8Al3BWPUnndM8ciJ4uQb/view?usp=drive_link

**Supplementary Video 1. Pheromone-exposed males move around more than control males.** Shown are traces of the paths taken by a representative pheromone-exposed male (top) and a control male (bottom). The green line shows previous spider positions.

https://drive.google.com/file/d/19zy0uv4SB4REzJz5HfUrzLIlNdGf1mOp/view?usp=drive_link

**Supplementary Video 2. Array presented to a male spider.** Video depicts an overlay between the retinal and stimulus positions. The spider first selects the cross, then views the rest of the stimuli in turn before disengaging with the array.

https://drive.google.com/file/d/1M5IWiNB0oRDfutYHYIlSxhW0CnqajhC1/view?usp=drive_link

**Supplementary Video 3. Array presented to a female spider.** Video depicts an overlay between the retinal and stimulus positions. The spider first selects the male conspecific image, then continues to inspect it with torsional retinal movements for repeated bouts.

https://drive.google.com/file/d/1eB10v2KCT7SYaJlEaqUcwBdIFVvPxOGI/view?usp=drive_link

**Supplementary Video 4. Spider keeps inspecting the initial stimulus.** Video depicts an overlay between the retinal and stimulus positions. The male spider ignores the distractor and keeps inspecting the initial stimulus.

https://drive.google.com/file/d/1qVdisCHr_AVGpvAukYkpj-LqfRu8pnwF/view?usp=drive_link

**Supplementary Video 5. Spider redirects its gaze to the distractor stimulus.** Video depicts an overlay between the retinal and stimulus positions. The male spider disengages with the initial stimulus and redirects its gaze to the distractor.

